# Detecting Random Mutations in 16S rRNA Sequences

**DOI:** 10.64898/2026.09.24.753821

**Authors:** Rain Haworth, Seth Commichaux, Mihai Pop

**Affiliations:** Department of Computer Science, University of Maryland, College Park, MD 20740; International Flavors and Fragrances, Wilmington, DE 19803

## Abstract

**Motivation:** High-throughput sequencing technologies have driven rapid growth of biological sequence databases. Public repositories must therefore rely on automated computational heuristics to screen submitted sequences for errors and low quality. For example, SILVA SSU Ref, which exploits the conserved nature of 16S and 18S rRNA sequences, applies strict algorithmic quality controls yet still accepts sequences with up to 30% of their nucleotides deviating from any previously accepted sequence. This permissiveness creates opportunities for the admission of modified sequences, such as biologically plausible sequences generated by DNA foundation models. The vulnerability of public sequence databases to becoming polluted or poisoned with modified sequences necessitates the development of methods to detect such sequences.

**Results:** We present the first investigation, to our knowledge, of the detectability of modified sequences. We consider simple computationally-modified 16S rRNA sequences that pass the quality control inclusion criteria of the SILVA database, which we generate via random substitutions. We present classifiers that can distinguish such modified sequences from natural 16S rRNA using conserved motifs. Our best classifier achieves over 90% sensitivity and specificity on our testing set when using a 5% artificial mutation rate. One feature used in our classifiers, gapped k-mers constructed from universally conserved nucleotides, was conserved across all three domains of life despite relying on exact matches to patterns found in *E. coli*, advancing our understanding of conserved grammatical structure in small subunit rRNA sequences.

**Availability and Implementation:** Our source code is available at https://github.com/rainhaworth/16S-Mutation-Classifiers.

## 1 Introduction

Since their inception in 1965, biological sequence databases have driven research progress in many fields, including molecular biology, medicine, agriculture, biotechnology, and microbial ecology [3, 4, 19]. Many biological sequence databases have seen continued rapid growth in sequence submissions [6], thus automated error detection tools and quality control heuristics have become the only practical way to reject undesired sequences. However, our poor understanding of the grammar of biological sequences has hindered the development of such heuristics, and many databases implement few quality controls despite accepting submissions from a vast range of sources. These databases are therefore vulnerable to pollution and poisoning by undesirable sequences, which may be low-quality, computationally generated, falsified, or engineered. The pollution of public databases with incorrect sequences can severely reduce the accuracy of the many computational analyses and tools that rely on public data sources.

We previously showed that the performance of the Ribosomal Database Project’s Bayesian taxonomic classifier can be degraded by adding a small number of computationally-generated sequences to its training data [9]. This classifier attempts to assign a genus label to 16S ribosomal RNA (rRNA) sequences. In one instance, a single computationally-generated sequence changed the predicted genus of 85% of sequences previously classified as *Dialister* to *Allisonella*, nearly erasing the presence of an entire genus from the analysis of a real stool microbiome dataset.

With the rapid development of large DNA foundation models like Evo 2 [2], it is becoming easier to computationally generate sequences that appear biologically plausible, thus it is increasingly likely that such sequences will be submitted to databases. Biological sequence databases therefore face a fundamental challenge: developing scalable computational methods that differentiate these from naturally occurring sequences.

To assess whether such methods can be developed, we focus on the SILVA rRNA sequence database [15, 16] because it uses well-defined computational quality controls to vet new sequences for admission. To admit a candidate sequence to the database, SILVA primarily considers the aggregate “sequence quality” score and three “alignment quality” metrics for each sequence. The sequence quality score aggregates the fraction of ambiguous base calls, homopolymers longer than four nucleotides, and regions with high identity to a known vector sequence. For alignment quality, SILVA places thresholds on the alignment score, base pair score, and alignment identity reported by SINA [14], an incremental alignment tool designed for SILVA. Throughout the database’s iterative construction process, each candidate sequence must therefore align to at least one sequence in the current database with some minimum identity threshold. In SSU Ref, the most stringently curated dataset containing 16S rRNA, this identity threshold is 70% as of release 111 [16].

Beyond database considerations, 16S rRNA sequences are highly conserved relative to other biological sequences and possess a comparatively well-characterized sequence grammar. This has prompted multiple efforts to model 16S rRNA sequences and their secondary structure using stochastic context-free grammars (SCFGs) [10, 17]. More recently, 140 nucleotides were identified as universally conserved across an alignment of 1,961 taxonomically diverse 16S rRNA sequences (Figure 1), suggesting the presence of core structural components essential for prokaryotic viability [11].

**Figure 1:**
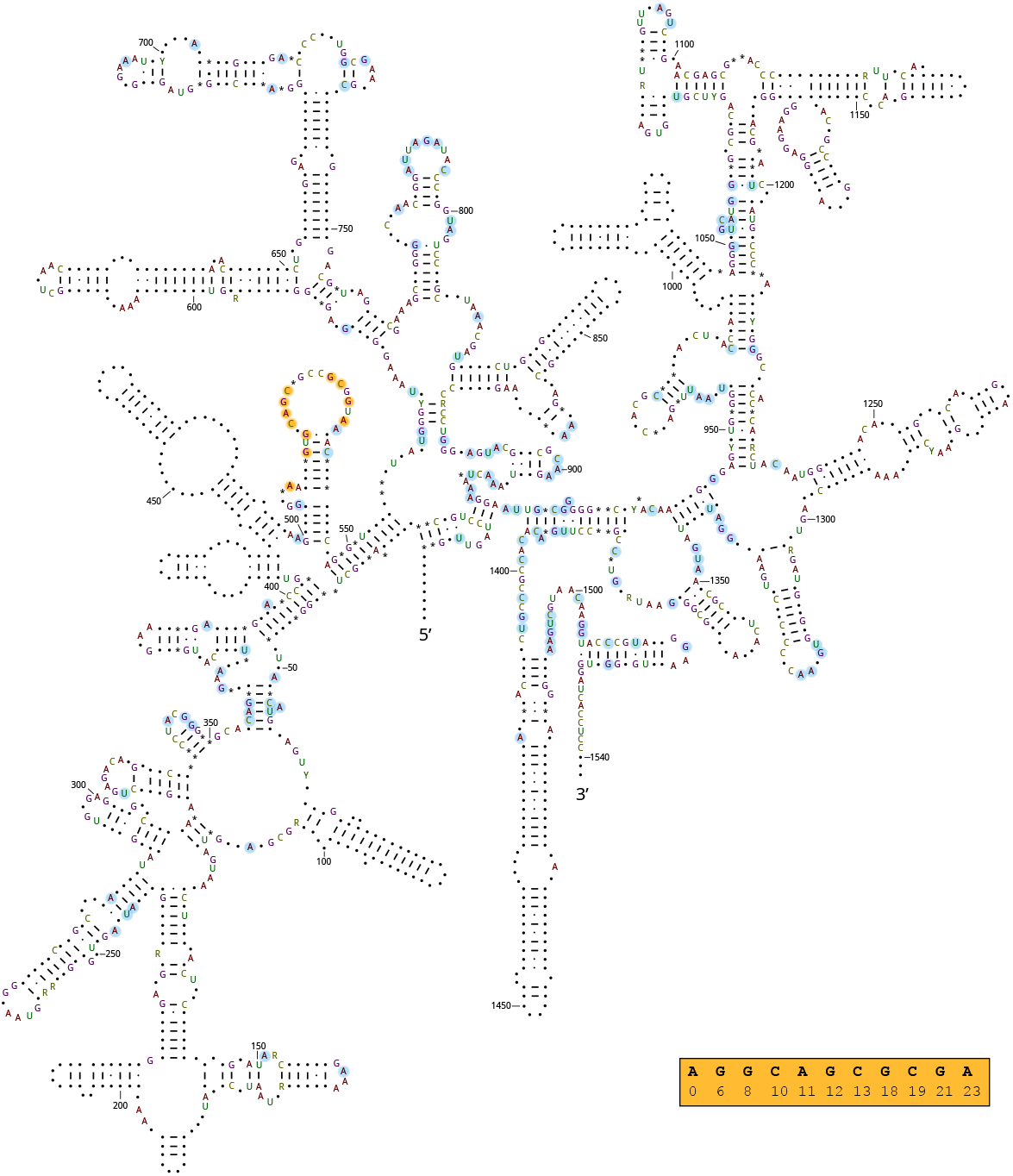
Secondary structure of 16S rRNA in *E. coli*, adapted from a 1987 illustration by C. R. Woese [23]. Nucleotides represented by a letter, rather than a symbol, are either heavily conserved or useful for distinguishing Bacteria from Archaea. Universally conserved nucleotides in 16S rRNA, identified by Noller et al. in 2022 [11], are highlighted in blue or orange. The orange box in the lower right contains an example gapped k-mer motif constructed from the sequence of nucleotides highlighted in orange (see Section 2.3.2).

Here, we assess whether this conservation and underlying grammatical structure can be leveraged to distinguish natural, full-length 16S rRNA sequences from computationally modified ones. By constructing modified sequence datasets that pass SILVA’s inclusion criteria, we test if additional sequence features can reliably discriminate biologically valid sequences from computationally generated ones. To simplify the problem setting, we restrict our analysis to modified sequences generated via random, independent substitutions at a fixed mutation rate, mimicking potential sequencing or experimental errors. Importantly, introducing random mutations is the least refined approach a hypothetical attacker may use if they tried to poison a biological database. This setting therefore provides an upper bound on the ability of computational tools to detect modified sequences.

## 2 Methods

We conducted a series of experiments with the goal of identifying individual features that can distinguish natural 16S rRNA sequences from randomly mutated ones. These experiments constitute a feature extraction phase, a training phase, and a testing phase. Datasets used in the testing phase are not seen during any other phase.

Formally, given a set *S* of known 16S rRNA sequences and another set 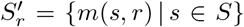, where *m*(*s, r*) mutates a fixed proportion *r ∈* (0, 1] of nucleotides in *s* to another random nucleotide, our objective was to build a classifier that correctly labels sequences as belonging to *S* and 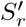. We considered each *s ∈ S* “positive” and each 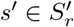 “negative.” An optimal classifier would produce only true positives and true negatives for any valid *r* and would generalize this property to any set of unseen 16S rRNA sequences *U* and its corresponding 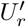.

In practice, we do not expect an optimal classifier to exist. Particularly at low values of *r*, artificial mutations may coincidentally fall within non-conserved regions and produce a mutated sequence that is indistinguishable from a natural sequence. With this limitation in mind, we evaluated classifiers across three mutation rates: *r* = 0.01, *r* = 0.05, and *r* = 0.1. These mutation rates are relevant for biological analysis of 16S rRNA, where 95% and 99% identity thresholds are commonly used to classify isolate bacteria genera and species [7, 18]. Additionally, 1% approaches a feasible error rate in properly collected sequence data. The final 10% mutation rate sits significantly above these thresholds, along with typical error rates for modern sequencing technologies, but well within the identity threshold used to reject candidate sequences from the SILVA database (see Section **??**).

### 2.1 Datasets

We conducted a series of experiments aiming to identify viable methods to distinguish natural 16S rRNA sequences from randomly mutated ones. Formally, given a set *S* of known 16S rRNA sequences and another set 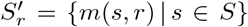, where *m*(*s, r*) mutates a fixed proportion *r ∈* (0, 1] of nucleotides in *s* to another random nucleotide, our objective was to build a classifier that correctly labels sequences as belonging to *S* and 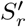. We considered each *s ∈ S* “positive” and each 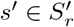 “negative.” An optimal classifier would produce only true positives and true negatives for any valid *r* and would generalize this property to any set of unseen 16S rRNA sequences *U* and its corresponding 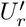.

In practice, we do not expect an optimal classifier to exist. At low values of *r*, artificial mutations may coincidentally fall within non-conserved regions of 16S rRNA genes and produce a modified sequence that is indistinguishable from a natural sequence. With this limitation in mind, we trained and tested classifiers across three mutation rates: *r* = 0.01, *r* = 0.05, and *r* = 0.1. These mutation rates are relevant for biological analysis of 16S rRNA, where 95% and 99% identity thresholds are commonly used to classify bacterial genera and species [7, 18]. Additionally, 1% approaches a feasible error rate in properly collected sequence data. The final 10% mutation rate sits above both these thresholds and typical error rates for modern sequencing technologies, but well within the identity threshold used to reject candidate sequences from the SILVA database.

Throughout this section, we sometimes use sequence data that are represented as DNA strings. In all such cases, we convert these strings to the RNA alphabet by replacing each T with U before any further processing.

### 2.2 Datasets

We constructed multiple training and testing datasets for evaluation. All rRNA sequences included in these datasets were found in SILVA [16]. We included two false positive testing sets because we were interested in determining whether our methods would incorrectly identify natural, non-16S nucleic acid sequences as valid 16S sequences. We therefore did not mutate these sequences and treated each natural sequence as a negative sample.

1. Three **SSU training sets** were constructed from all sequences from SILVA SSU Ref NR 99 excluding Eukaryotic 18S rRNA, sequences from MAGs, and sequences with chimerism or anomalies reported by Pintail [1]. This additional filtering retains 282,395 of the original 510,495 sequences in SSU Ref NR 99. Each natural sequence is used as a positive sample, while modified sequences mutated at a fixed *r ∈ {*0.01, 0.05, 0.1} are used as negative samples; each final dataset therefore contains 564,790 sequences.
2. Three **SSU testing sets** were constructed from all sequences in SILVA SSU Ref. For classifier testing, we exclude sequences found in our ground truth training set and Eukaryotic 18S rRNA, retaining 1,769,615 of the original 2,224,690 sequences. Positive and negative samples are generated by the same procedure used for our SSU training sets with the same values of *r*.
3. One **large subunit (LSU) false positive testing set** contained all 23S and 28S rRNA sequences in SILVA LSU Ref.
4. One **non-RNA false positive testing set** contained 115,561 bacterial protein-coding genes collected from 33 diverse NCBI RefSeq bacterial genomes in the kingdoms *Bacillati* and *Pseudomonadati*. Genome metadata are listed on GitHub in the file `ncbi_dataset.tsv`.

Additionally, in Section 3.3, we investigate the robustness of the most successful features across all sequences in SSU Ref, including Eukaryotic 18S, grouped by taxonomy.

### 2.3 Candidate distinguishing sequence features

To construct successful classifiers, we must identify attributes of sequences whose distributions differ significantly between natural and modified sequences. We designed 30 features that we expect to have this property and computed them separately for each sequence.

#### 2.3.1 Statistical features

Features in this category do not require any prior knowledge of 16S rRNA to compute. Each can be computed in *O*(*n*) time, where *n* is the number of nucleotides in the sequences being analyzed.

1. **Nucleotide statistics:** the proportions of each nucleotide (A, C, G, U); all ratios of one nucleotide count to another, e.g., A/C, C/G; and GC-content, the fraction of the sequence containing either a G or a C.
2. **Homopolymers**, defined here as the number of separate, contiguous subsequences containing four or more consecutive identical nucleotides, e.g., AAAA. We consider both the number of homopolymers of each nucleotide and the ratio of G homopolymers to total homopolymers. We introduced the latter feature after we observed a strong bias toward G homopolymers in natural 16S rRNA gene sequences.
3. **K-mer statistics** constructed from the number of occurrences of each k-mer for *k ∈* [1, 3, 5, 7, 9]. Our first statistic is **k-mer parity**, which we define as the proportion of k-mers where both the k-mer itself and its reverse complement are present in the sequence. We expect k-mer parity to decrease in modified sequences due to the degradation of natural complementarity in the secondary structure of 16S rRNA. Our second statistic is the **Shannon entropy** *H*(*X*) = − ∑_*x ∈X*_ *f* (*x*) log *f* (*x*) of all non-zero k-mer frequencies *f* (*x*) = *x/N*. Here we expect the addition of random noise to produce a more diverse k-mer composition and therefore increase the Shannon entropy.

#### 2.3.2 Motifs in 16S rRNA

We consider three types of motifs we expect to occur widely across 16S rRNA. For each feature in this category, we construct a fixed set of motifs *M* then, for each sequence *s*, count the number of unique motifs *{m ∈ M* | *m ∈ s}* found. These features can be computed in *O*(*mn*) time.

1. **Universal primers** are DNA sequences capable of initiating PCR amplification of 16S rRNA in a wide range of microbial taxa [22]. To achieve this, they must match highly conserved regions found exclusively in 16S rRNA sequences. We collected a list of 14 universal primers: 8F/FD1 and 2 variants of 1492R/RP1 from [22]; 27F, 337F, and 518R from [13]; 336R, 907R, and 928F from [21]; 515F and 806R from [5]; 785F from [12]; and 1100F and 1100R from [20].
2. We identified 295 **frequent k-mers** of length *k* = 11 occurring in at least 50% of natural sequences in our training set.
3. We constructed **gapped k-mers** of length *k* = 11 from consecutive positions of universally conserved nucleotides in *E. coli*, as shown in Figure 1. For each nucleotide *n*_*i*_ at position *p*_*i*_, the corresponding gapped k-mer contains *k* pairs 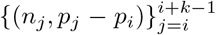 representing the relative offsets of each nucleotide from *p*_*i*_. To limit our gapped k-mer set to those modeling relatively local structures, we excluded those whose width *p*_*i*+*k™*1_*™ p*_*i*_ exceeded 100, producing 72 accepted k-mers out of 130 candidates. These gapped k-mers occur in 15% to 95% of natural sequences in our training set.

For both k-mer motifs, we selected *k* = 11 following initial experiments in which lower values of *k* produced motifs that were too common in modified sequences, while higher values of *k* produced motifs that were too scarce in natural sequences.

#### 2.3.3 Predicted secondary structure

Natural 16S rRNA must fold into a specific structure (see Figure 1) to function properly in an organism’s ribosomes. With the intuition that random mutations would disrupt this structure and, therefore, would be easily discoverable, we directly predicted RNA structure with ViennaRNA’s RNAFold [8], which runs in *O*(*n*^3^) time. We use the reported Gibbs free energy and the number of predicted helices as features.

### 2.4 SILVA inclusion screening of modified sequences

To assess whether the modified sequences would pass SILVA inclusion criteria, we aligned each modified training sequence to the SILVA SSU Ref NR 99 ARB reference using SINA. Sequences achieving an alignment quality of *≥* 50, a base pair score of *≥* 30, and with *≥* 70% identity to a representative SILVA database sequence were considered to pass inclusion. We do not apply SILVA sequence quality scores, as random mutations of SILVA reference sequences will not introduce ambiguous bases or vector contamination and are unlikely to generate long homopolymers.

### 2.5 Feature and classifier evaluation procedure

We first evaluated each individual candidate feature by both training and testing a logistic regression classifier on our training set with an 80*/*20 train/test split. Here, we considered only the highest mutation rate, i.e., *r* = 0.1. Features whose classifiers achieved at least 80% sensitivity and specificity advanced to the next round of experiments.

Accepted features had their logistic regression classifiers retrained on the full training set at each mutation rate. These features were also combined to train decision tree classifiers. To investigate the relationship between performance and model complexity, we trained decision tree classifiers with a range of maximum depth restrictions: 3, 5, 10, and unlimited. Each classifier in this final set was evaluated on all three test sets.

## 3 Results

### 3.1 SILVA inclusion screening of modified sequences

To determine whether SILVA’s filtering based on SINA alignment metrics would reject modified 16S rRNA sequences, we used the sequences with the highest mutation rate, *r* = 0.1. None of the modified sequences fell within SILVA’s rejection threshold for alignment quality, base pair score, or alignment identity metrics (Table 1). This confirmed that SINA would not filter out these modified sequences.

**Table 1:** SINA metrics when aligning the modified sequences to the SSU Ref NR 99 dataset. Note that base pair score can take on values above 100, while alignment score and alignment identity are normalized to [0, 100].

| Metric | Pass Rate | Mean | Min. | Thresh. |
| --- | --- | --- | --- | --- |
| Alignment score | 100% | 85.86 | 57 | 50 |
| Base pair score | 100% | 100.35 | 46 | 30 |
| Alignment identity | 100% | 89.85 | 71.01 | 70 |

### 3.2 Feature selection

As seen in Table 2, three features exceed our acceptance threshold of 80% sensitivity and specificity: universal primers, gapped k-mers, and frequent k-mers.

**Table 2:** Sensitivity and specificity of single-feature logistic regression classifiers on the training set at *r* = 0.1. Time indicates the total wall-clock time required to compute each feature for all sequences, excluding preprocessing and classifier operations. * - Only evaluated over 100 sequences due to long runtimes; runtime is reported as an estimate on the full dataset.

| Feature | Sens. | Spec. | Time (m) |
| --- | --- | --- | --- |
| A | 100.0 | 0.00 | } 0.7 |
| C | 56.46 | 52.69 |  |
| G | 63.24 | 63.09 |  |
| U | 60.03 | 59.70 |  |
| A/C | 52.77 | 53.27 |  |
| A/G | 57.54 | 55.11 |  |
| A/U | 60.10 | 61.76 |  |
| C/G | 69.27 | 65.95 |  |
| C/U | 49.34 | 57.66 |  |
| G/U | 60.47 | 64.30 |  |
| GC-content | 54.77 | 56.16 |  |
| HomopolymerA | 58.75 | 61.04 | } 2.9 |
| HomopolymerC | 52.24 | 58.26 |  |
| HomopolymerG | 61.52 | 56.09 |  |
| HomopolymerU | 52.66 | 54.49 |  |
| G homopolymer bias | 64.78 | 63.13 |  |
| Parity, $k = 1$ | 71.20 | 69.70 | } 23.7 |
| Parity, $k = 3$ | 71.33 | 68.72 | |
| Parity, $k = 5$ | 65.44 | 67.19 | |
| Parity, $k = 7$ | 63.91 | 61.34 | |
| Parity, $k = 9$ | 65.08 | 52.59 | |
| Entropy, $k = 1$ | 56.39 | 71.04 | |
| Entropy, $k = 3$ | 57.32 | 70.70 | |
| Entropy, $k = 5$ | 53.57 | 68.48 | |
| Entropy, $k = 7$ | 42.46 | 60.37 | |
| Entropy, $k = 9$ | 100.0 | 0.00 | |
| Universal primers | **87.65** | **98.50** | 1.4 |
| Frequent k-mers | **91.60** | **99.16** | 10.6 |
| Gapped k-mers | **93.46** | **94.52** | 9.3 |
| Gibbs free energy* | 42.86 | 47.37 | } 36,029 |
| Helices* | 52.38 | 52.63 |  |

Notably, our most computationally expensive features, based on secondary structure predictions, had poor classifier performance and an estimated runtime three orders of magnitude greater than the next most expensive feature. We attribute these observations primarily to the inherent challenges of predicting secondary structure. RNAfold achieves its *O*(*n*^3^) time complexity by ignoring pseudoknots and ranking structures by Gibbs free energy [8], which frequently produces multiple divergent candidates that may all deviate significantly from the true secondary structure of the sequence. Due to these limitations and the poor classifier performance, we did not further investigate features based on predicted structure.

### 3.3 Performance of finalized classifiers

We evaluated the three successful features identified in the previous section on our three test tasks: distinguishing unseen natural 16S rRNA sequences from unseen modified sequences, searching LSU Ref for false positives, and searching DNA genes for false positives. In Table 3, we examine each single feature classifier and each decision tree. We select maximum decision tree depth restrictions *d ∈ {*3, 5, 10*}* to assess the variation of performance with model complexity.

**Table 3:**
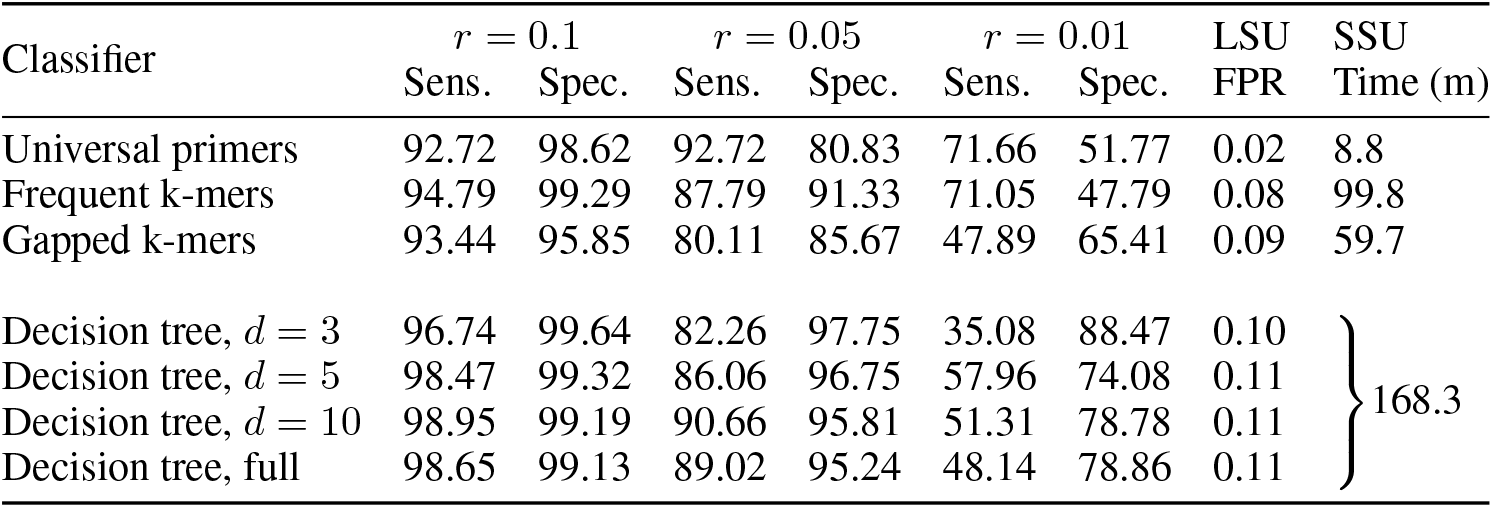
Performance of the final set of classifiers on 16S rRNA sequences and LSU rRNA sequences in our testing sets. Sensitivity, specificity, and false positive rate (FPR) are reported as percentages. Runtime was computed on all sequences in SSU Ref. We exclude DNA genes from this table because none of our classifiers found any false positives.

Across all classifiers, we observe low false positive rates on LSU Ref and no false positives on DNA genes. All classifiers achieve at least 80% sensitivity and specificity at *r* = 0.1 and *r* = 0.05, while none achieve 70% sensitivity and specificity at *r* = 0.01. The performance of the decision trees increases with model complexity up to *d* = 10, but the full model sees a small drop in performance across all mutation rates.

### 3.4 Domain-stratified analysis

The results shown in previous sections are shaped by the taxonomic imbalance of SSU Ref, which is dominated by Bacteria, constituting approximately 92% of training sequences and 97% of testing sequences This particularly skews the results in Table 3 in favor of features that perform well on Bacteria. As such, we assess the strength of our methods across Bacteria, the majority class; Archaea, the minority class; and Eukaryota, an unseen class. We begin by testing our existing classifiers on each domain of sequences in SSU Ref at *r* = 0.1, as shown in Table 4.

**Table 4:**
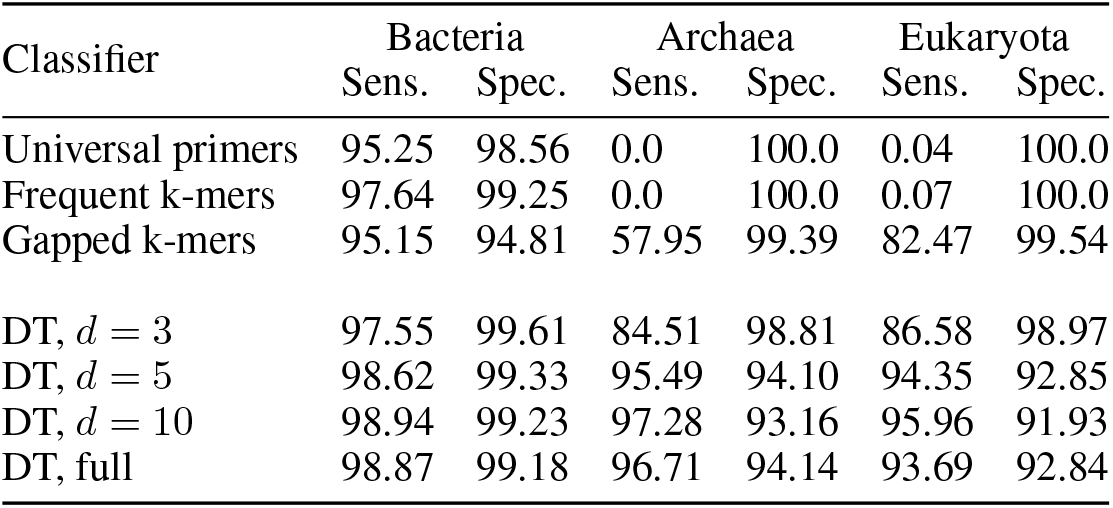
Performance of each classifier on all sequences in SSU Ref at *r* = 0.1, grouped by domain. DT refers to the decision tree classifiers.

This analysis revealed that, among our three individual features, only gapped k-mers can fit a single threshold to (primarily) Bacteria and generalize to both Archaea and Eukaryota. The other single feature classifiers outperform gapped k-mers on Bacteria, but they fail to identify a single natural sequence within Archaea and only produce true positives for a small minority (*<* 0.1%) of natural sequences within Eukaryota. Nonetheless, the decision trees combining information from all three features perform substantially better than gapped k-mers alone on Archaea and Eukaryota. The strong performance on unseen Eukaryota indicates limited overfitting at least up to a maximum depth of *d* = 10.

We then examined the prevalence of each individual feature in natural and modified 16S and 18S rRNA sequences grouped at the family level (Figure 2). To minimize visual noise, we depicted only the mean and standard deviation within each phylogroup, sorted our results by mean, and sorted natural and modified sequences separately while enforcing strict domain separation. Informative features should have a high prevalence in all phylogroups with a large separation between natural and modified sequences.

**Figure 2:**
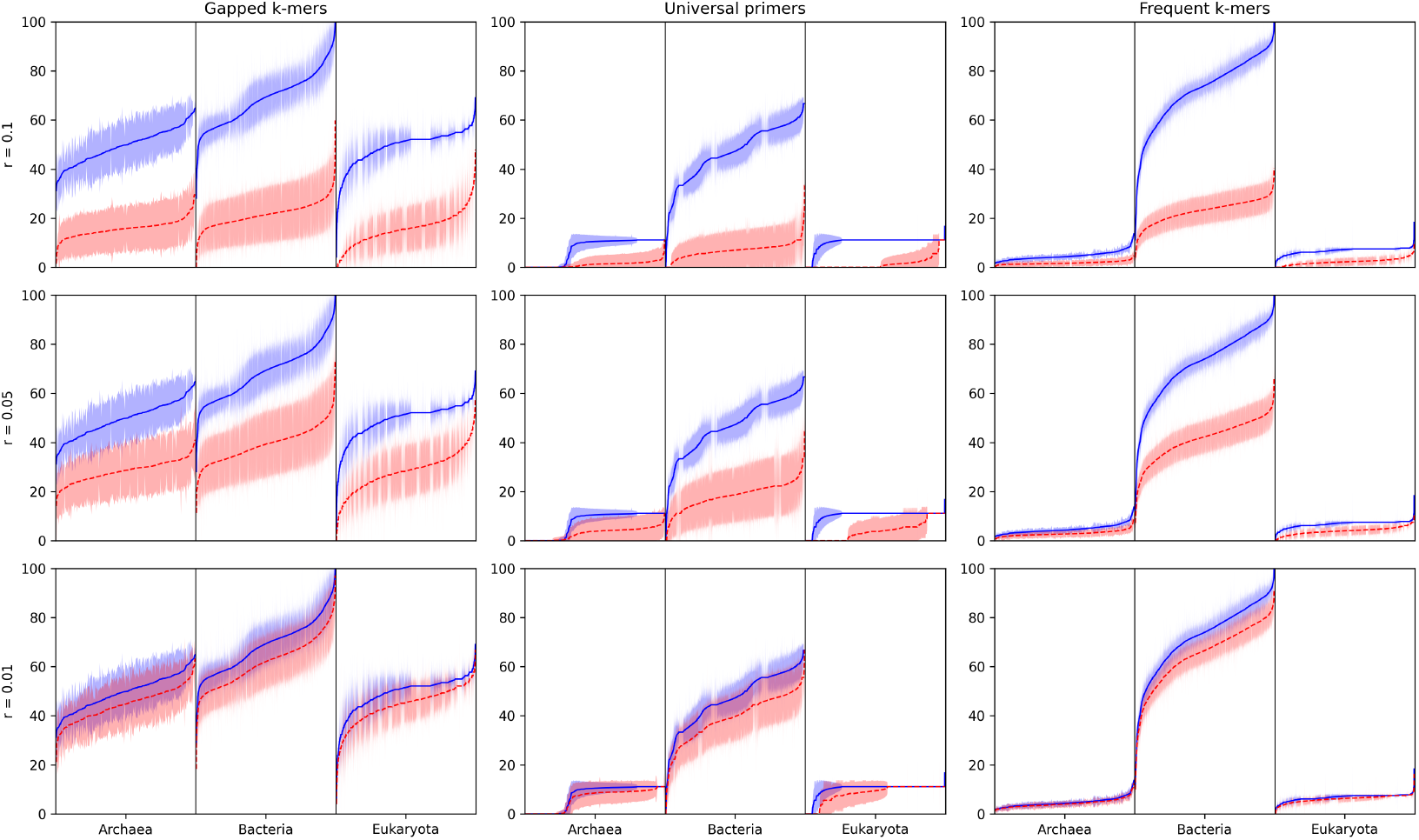
Percentage of each motif found in natural (solid blue lines) and modified (dashed red lines) SSU rRNA sequences, aggregated into taxonomic groups then sorted in ascending order. Each vertical slice depicts *µ ± σ* of the percentage of motifs found within one taxonomic group (e.g., family) listed in SSU Ref. This figure contains 275 taxonomic groups in Archaea, 4,853 in Bacteria, and 6,158 in Eukaryota. As a result of our sorting method, the order of taxa varies both across graphs and between natural and mutated sequences in the same graph. A large gap between the solid and dashed line (and associated error bands) suggests that a simple classifier could distinguish natural sequences from mutated ones.

The results of this experiment support our suggestion that gapped k-mers are most suitable for building a “one size fits all” classifier for 16S and 18S SSU rRNA, with similar separation between real and mutated sequences and relatively mild variance across taxa. Conversely, the universal primer and frequent k-mer features are universally far less abundant in Archaea and Eukaryota than in Bacteria, with much weaker separation between real and mutated sequences in these domains. This suggests that classifiers using the universal primer and frequent k-mer features might be improved by incorporating domain awareness, for example by using domain-specific thresholds or motif sets.

### 3.5 Runtime

Our Python implementation computed all three features at a speed of over 1.5 million sequences per hour on a single CPU core. For comparison, the alignment of 282,395 sequences against the SSU Ref NR 99 database with SINA had a runtime over 16 hours, equivalent to a speed of approximately 17,000 sequences per hour.

## 4 Discussion

We have identified three 16S rRNA motif features that can detect modified sequences that would be accepted by SILVA, according to metrics reported by SINA. We find that sequences modified at a 10% or greater induced mutation rate can be detected with reasonable accuracy (*≥* 90%) using any individual feature or with high accuracy (*≥* 98%) using a decision tree. At lower mutation rates, we only advise the use of a decision tree. Each classifier’s performance represents a tradeoff between acceptance of natural sequences (sensitivity) and rejection of mutated sequences (specificity) for any given *r*. At a 1% mutation rate, none of our classifiers achieved high performance, likely because this mutation rate is unlikely to destroy enough conserved local structures to be distinguishable from natural mutations.

The success of gapped k-mer motifs across all domains of life builds upon recent discoveries about SSU rRNA grammar [11]. First, it confirms that many of these nucleotides remain conserved in a much larger dataset, though not as universally as suggested by the original paper. Second, the success of exact matching against the relative nucleotide positions in *E. coli* demonstrates that many local structures are heavily conserved with identical gap lengths between structurally relevant nucleotides. However, our investigation of these gapped k-mers is somewhat shallow, examining them only in aggregate and at a single value of *k*. Future work should examine variation across taxa and potential functional roles for each individual k-mer; search for the presence of slight variations in shape; examine different k-mer lengths; and consider merging or separating k-mers into groupings that more accurately reflect conserved structural units.

The high performance of universal primers and frequent k-mers in Bacteria, despite their failure to generalize to Archaea and Eukaryota, is somewhat explainable. Many universal primers were designed for conserved sites in Bacteria which are often less conserved in or entirely absent from Archaea or Eukaryota. Frequent k-mers were biased toward Bacteria because they constitute 92% of sequences in our training set; a “domain-aware” approach accepting k-mers either frequent in only one domain or frequent across multiple domains may be able to perform well on multiple domains while maintaining generalization to unseen sequences.

Although this case study was limited to detecting naively modified sequences, it highlights the vulnerability of public sequence databases to pollution or poisoning by modified sequences. Future work should continue to investigate this risk, particularly the detection of more sophisticated modifications, including those that can now be generated by DNA foundation models such as Evo 2 [2]. Central to this effort is a deeper understanding of biological sequence grammar, enabling biologically functional sequences to be distinguished from the vast space of possible sequence combinations.

## 5 Data availability statement

The data underlying this article are available in SILVA at

- https://dx.doi.org/10.82364/138.2/SSU/Ref/FASTA/unaligned,
- https://dx.doi.org/10.82364/138.2/SSU/Ref-NR99/FASTA/unaligned, and
- https://dx.doi.org/10.82364/138.2/LSU/Ref/FASTA/unaligned

and in NCBI RefSeq at accessions listed in the file `ncbi_dataset.tsv` provided in our GitHub repository.

## 6 Acknowledgments

Research reported in this publication was supported by the National Library Of Medicine of the National Institutes of Health under Award Number R01LM014698. The content is solely the responsibility of the authors and does not necessarily represent the official views of the National Institutes of Health.

## References

[1] Kevin E. Ashelford et al. “At Least 1 in 20 16S rRNA Sequence Records Currently Held in Public Repositories Is Estimated To Contain Substantial Anomalies”. In: Applied and Environmental Microbiology 71.12 (Dec. 2005), pp. 7724–7736. DOI: 10.1128/AEM.71.12.7724-7736.2005. URL: https://journals.asm.org/doi/10.1128/aem.71.12.7724-7736.2005 (visited on 05/27/2025).

[2] Garyk Brixi et al. “Genome modelling and design across all domains of life with Evo 2”. In: Nature (2026), pp. 1–13.

[3] Christian Burks et al. “GenBank”. In: Nucleic Acids Research 20.Suppl (May 1992), pp. 2065–2069. ISSN: 0305-1048. DOI: 10.1093/nar/20.suppl.2065. URL: https://doi.org/10.1093/nar/20.suppl.2065 (visited on 09/26/2025).

[4] Jeff Gauthier et al. “A brief history of bioinformatics”. In: Briefings in Bioinformatics 20.6 (Nov. 2019), pp. 1981–1996. ISSN: 1477-4054. DOI: 10.1093/bib/bby063. URL: https://doi.org/10.1093/bib/bby063 (visited on 10/13/2025).

[5] Jack A Gilbert et al. “The Earth Microbiome Project: Meeting report of the “1st EMP meeting on sample selection and acquisition” at Argonne National Laboratory October 6th 2010”. In: Standards in genomic sciences 3.3 (2010), pp. 249–253.

[6] Kenneth Katz et al. “The Sequence Read Archive: a decade more of explosive growth”. In: Nucleic acids research 50.D1 (2022), pp. D387–D390.

[7] Mincheol Kim et al. “Towards a taxonomic coherence between average nucleotide identity and 16S rRNA gene sequence similarity for species demarcation of prokaryotes”. In: International Journal of Systematic and Evolutionary Microbiology 64.Pt_2 (2014), pp. 346–351. ISSN: 1466-5034. DOI: 10.1099/ijs.0.059774-0. URL: https://www.microbiologyresearch.org/content/journal/ijsem/10.1099/ijs.0.059774-0 (visited on 09/22/2025).

[8] Ronny Lorenz et al. “ViennaRNA Package 2.0”. en. In: Algorithms for Molecular Biology 6.1 (Nov. 2011), p. 26. ISSN: 1748-7188. DOI: 10.1186/1748-7188-6-26. URL: https://doi.org/10.1186/1748-7188-6-26 (visited on 09/18/2025).

[9] Harihara Subrahmaniam Muralidharan, Noam Y. Fox, and Mihai Pop. “The impact of transitive annotation on the training of taxonomic classifiers”. English. In: Frontiers in Microbiology 14 (Jan. 2024). ISSN: 1664-302X. DOI: 10.3389/fmicb.2023.1240957. URL: https://www.frontiersin.org/journals/microbiology/articles/10.3389/fmicb.2023.1240957/full (visited on 04/07/2025).

[10] Eric P. Nawrocki, Diana L. Kolbe, and Sean R. Eddy. “Infernal 1.0: inference of RNA alignments”. In: Bioinformatics 25.10 (May 2009), pp. 1335–1337. ISSN: 1367-4803. DOI: 10.1093/bioinformatics/btp157. URL: https://doi.org/10.1093/bioinformatics/btp157 (visited on 09/18/2025).

[11] Harry F. Noller, John Paul Donohue, and Robin R. Gutell. “The universally conserved nucleotides of the small subunit ribosomal RNAs”. In: RNA 28.5 (May 2022), pp. 623–644. ISSN: 1355-8382. DOI: 10.1261/rna.079019.121. URL: https://www.ncbi.nlm.nih.gov/pmc/articles/PMC9014874/ (visited on 04/02/2025).

[12] AR Özok et al. “Ecology of the microbiome of the infected root canal system: a comparison between apical and coronal root segments”. In: International endodontic journal 45.6 (2012), pp. 530–541.

[13] Changwoo Park et al. “Comparison of 16S rRNA gene based microbial profiling using five next-generation sequencers and various primers”. In: Frontiers in Microbiology 12 (2021), p. 715500.

[14] Elmar Pruesse, Jörg Peplies, and Frank Oliver Glöckner. “SINA: Accurate high-throughput multiple sequence alignment of ribosomal RNA genes”. In: Bioinformatics 28.14 (July 2012), pp. 1823–1829. ISSN: 1367-4803. DOI: 10.1093/bioinformatics/bts252. URL: https://doi.org/10.1093/bioinformatics/bts252 (visited on 05/07/2025).

[15] Elmar Pruesse et al. “SILVA: a comprehensive online resource for quality checked and aligned ribosomal RNA sequence data compatible with ARB”. In: Nucleic Acids Research 35.21 (Dec. 2007), pp. 7188–7196. ISSN: 0305-1048. DOI: 10.1093/nar/gkm864. URL: https://doi.org/10.1093/nar/gkm864 (visited on 05/07/2025).

[16] Christian Quast et al. “The SILVA ribosomal RNA gene database project: improved data processing and web-based tools”. In: Nucleic Acids Research 41 (D1 Jan. 1, 2013), pp. D590–D596. ISSN: 0305-1048. DOI: 10.1093/nar/gks1219. URL: https://doi.org/10.1093/nar/gks1219 (visited on 04/30/2025).

[17] Elena Rivas and Sean R. Eddy. “The language of RNA: a formal grammar that includes pseudoknots”. In: Bioinformatics 16.4 (Apr. 2000), pp. 334–340. ISSN: 1367-4803. DOI: 10.1093/bioinformatics/16.4.334. URL: https://doi.org/10.1093/bioinformatics/16.4.334 (visited on 09/22/2025).

[18] Morgane Rossi-Tamisier et al. “Cautionary tale of using 16S rRNA gene sequence similarity values in identification of human-associated bacterial species”. In: International Journal of Systematic and Evolutionary Microbiology 65.Pt_6 (2015), pp. 1929–1934. ISSN: 1466-5034. DOI: 10.1099/ijs.0.000161. URL: https://www.microbiologyresearch.org/content/journal/ijsem/10.1099/ijs.0.000161 (visited on 09/18/2025).

[19] Guenter Stoesser et al. “The EMBL Nucleotide Sequence Database”. In: Nucleic Acids Research 27.1 (Jan. 1999), pp. 18–24. ISSN: 0305-1048. DOI: 10.1093/nar/27.1.18. URL: https://doi.org/10.1093/nar/27.1.18 (visited on 09/18/2025).

[20] Sean Turner et al. “Investigating deep phylogenetic relationships among cyanobacteria and plastids by small subunit rRNA sequence analysis 1”. In: Journal of Eukaryotic Microbiology 46.4 (1999), pp. 327–338.

[21] Stefan Weidner, Walter Arnold, and Alfred Puhler. “Diversity of uncultured microorganisms associated with the seagrass Halophila stipulacea estimated by restriction fragment length polymorphism analysis of PCR-amplified 16S rRNA genes”. In: Applied and environmental microbiology 62.3 (1996), pp. 766–771.

[22] William G Weisburg et al. “16S ribosomal DNA amplification for phylogenetic study”. In: Journal of bacteriology 173.2 (1991), pp. 697–703.

[23] Carl R Woese. “Bacterial evolution”. In: Microbiological reviews 51.2 (1987), pp. 221–271.

